# PHB3 Regulates Lateral Root Primordia Formation via NO-mediated Degradation of AUX/IAAs

**DOI:** 10.1101/2021.10.28.466218

**Authors:** Shuna Li, Qingqing Li, Xiao Tian, Lijun Mu, Meiling Ji, Xiaoping Wang, Na Li, Fei Liu, Jing Shu, Nigel M. Crawford, Yong Wang

## Abstract

We previously showed that *PHB3* regulates auxin-stimulated lateral root (LR) formation; however, the underlying molecular mechanism is unknown. Here, we demonstrate that *PHB3* regulates LR development mainly through influencing lateral root primordia (LRP) initiation via affecting nitric oxide (NO) accumulation. The reduced LRP in *phb3* was largely rescued by exogenous NO donor SNAP treatment. The decreased NO accumulation in *phb3* caused a lower expression of *GATA23* and *LBD16* through inhibiting the degradation of IAA14/28. Overexpression of either *GATA23* or *LBD16* in *phb3* mutant background recovered the reduced LRP number phenotype. These results indicate that PHB3 regulates LRP initiation via NO-mediated auxin signaling through regulating the degradation of IAA14/28.

**Highlight:** PHB3 regulates IAA28 and IAA14 degradation by controlling NO accumulation, and thereby regulating LR founder cell identification and the transition to asymmetric division.

## Introduction

Root architecture is an important factor influencing crop yield. The plasticity of lateral roots (LRs) allows the root architecture to adjust rapidly to the rhizosphere environment, enhancing resistance to various stresses (De Smet *et al*., 2012). Signaling molecules, especially hormones and nitric oxide (NO), play important roles in LR development (Fukaki & Tasaka, 2009; Jin *et al*., 2011; Sun *et al*., 2015).

Auxin signaling is critical for LR development and is mediated by the auxin co-receptors TRANSPORT INHIBITOR RESPONSE1/AUXIN SIGNALING F-BOX (TIR1/AFB), the transcriptional repressors AUXIN/INDOLE-3-ACETIC ACID (AUX/IAA), and AUXIN RESPONSE FACTORs (ARFs) (Correa-Aragunde, Graziano, & Lamattina, 2004; Laskowski *et al*., 2008; Fukaki & Tasaka, 2009; Lavenus *et al*., 2013; Guseman *et al*., 2015). The auxin signaling cascade is activated by the binding of auxin to the F-box auxin receptor complexes SCF^TIR1/AFBs^ followed by ubiquitination of AUX/IAA proteins, which are then degraded by 26S proteasomes (Lavy & Estelle, 2016; Leyser, 2018). LR initiation involves the specification and asymmetric division of LR founder cells (De Rybel *et al*., 2010; Goh, Joi, Mimura, & Fukaki, 2012; Teixeira and Tusscher, 2019; Van Norman *et al*., 2013). Of special importance to LR initiation are IAA28 and IAA14. IAA28 plays a key role in specifying pericycle cells for development into LR founder cells prior to LR initiation (De Rybel *et al*., 2010), while IAA14 is involved in the asymmetric division of LR founder cells that occurs as part of LR initiation (Fukaki *et al*., 2005; Goh, Joi, Mimura, & Fukaki, 2012). IAA28 can interact with ARF5/6/7/8/19. When IAA28 is degraded, ARF5/6/7/8/19 are free to activate *GATA23* expression, resulting in the specification of LR founder cells (De Rybel *et al*., 2010; Guseman *et al*., 2015). Next, auxin induces the degradation of IAA14/SLR, allowing ARF7/19 to activate the expression of *LBD16*, which then promotes the asymmetric division required for the migration of two adjacent founder cell nuclei to the common cell wall (Fukaki *et al*., 2005; Goh, Joi, Mimura, & Fukaki, 2012). Then, the IAA14/SLR-ARF7/19 and IAA12/BDL-ARF5 modules regulate the first round of asymmetric cell division and initiate LR primordia (LRP) formation (Fukaki *et al*., 2002; Okushima *et al*., 2005; Lavenus *et al*., 2013).

The NO free radical has been reported to play important roles in a variety of intracellular signaling in plants (Sengoku *et al*., 2015). NO acts as a second messenger that coordinates with hormone signaling to regulate growth, development, and certain physiological processes (Zandonadi *et al*., 2010; Terrile *et al*., 2012; Arc *et al*., 2013). It has been found that auxin can promote NO accumulation and NO is involved in auxin-induced LR development in tomato (*Solanum lycopersicum*) and *Arabidopsis* (Correa-Aragunde, Graziano, & Lamattina, 2004; Kolbert *et al*., 2008). Research into the mechanisms by which NO regulates LR development has revealed that NO influences auxin signaling through S-nitrosylation of ASK1 and TIR1 (Terrile *et al*., 2012; lglesias *et al*., 2018). Nitrosylation of ASK1 promotes the SCF^TIR1/AFBs^ E3 ligase assembly (lglesias *et al*., 2018) and TIR1 S-nitrosylation can enhance the interaction of TIR1/AFB with AUX/IAA, and thereby promote the degradation of AUX/IAA proteins and the expression of auxin-induced genes (Terrile *et al*., 2012).

Prohibitins (PHBs), originally discovered as tumor□suppressors in mammalian cells, are pleiotropic scaffold proteins involved in many important cell functions, including survival, proliferation, metabolism, and mitochondrial dynamics (Wang *et al*., 2020) and exist widely in plants and animals. There are seven conserved *PHB* genes in *Arabidopsis*, including *PHB3*, which is mainly localized in mitochondria and involved in regulating plant development and stress responses (Van Aken *et al*., 2007; Wang *et al*., 2010; Seguel *et al*., 2018; Huang *et al*., 2019). Dr. Larsen’s group has demonstrated that PHB3 plays an important role in modulating ethylene signaling in *Arabidopsis* and that its mutant *eer3* shows reduced size of rosette leaves (Christians & Larsen, 2007). Dr. Holuigue’s group showed that PHB3 interacts with ISOCHORISMATE SYNTHASE1 (ICS1), an essential enzyme in salicylic acid (SA) biosynthesis, and that the *phb3* mutant exhibits reduced SA generation and impaired responses to stresses such as pathogen infection (Seguel *et al*., 2018).

In a previous study, we found that H_2_O_2_-induced NO accumulation was inhibited in *phb3* mutants (Wang *et al*., 2010). Furthermore, auxin-induced LR development was reduced in *phb3* mutants (Wang *et al*., 2010). *PHB3* is mainly expressed in actively dividing tissues such as root tips, stems, and LRs (Kong *et al*., 2018). When *Pro*AtPHB:GFP-GUS transgenic plants, carrying a *GFP-GUS* reporter gene attached to the *PHB3* promoter, were treated with auxin 1-naphthaleneacetic acid (NAA), *PHB3* promoter activity could be detected in pericycle cells and was induced by NAA before the asymmetric division of founder cells (Van Aken *et al*., 2007). These findings imply that PHB3 may be involved in auxin-induced LR initiation and may be a key factor connecting endogenous NO accumulation and auxin-induced LR development, although the underlying mechanism is unknown.

Here, we demonstrate that *PHB3* regulates LRP initiation by controlling endogenous NO accumulation in LRPs. Treatments with NO induced the expression of *GATA23* and *LBD16*. However, *phb3* mutants showed reduced levels of *GATA23* and *LBD16* expression, and this reduction was correlated with strong inhibition of IAA28 and IAA14 degradation. These findings suggest that PHB3 regulates NO-mediated degradation of IAA28 and IAA14 and affects LRP initiation by regulating the expression of *GATA23* and *LBD16*.

## Materials and Methods

### Plant materials and *Arabidopsis* transformation

The *Arabidopsis* mutant lines *gata23* (De Rybel *et al*., 2010) and *lbd16* (SALK_095791) and transgenic line *pLBD16::GUS* were provided by colleagues and *phb3-3/ back26-2* mutant was generated as described by Wang *et al*. (2010). The *phb3ko* (SALK_020707) mutant was ordered from the Arabidopsis Biological Resource Center (ABRC) and verified by RT-PCR.

The *pPHB3::GUS* constructs were generated by introducing the histochemically detectable *β*-glucuronidase (*GUS*) gene into the pPZP211 vector and inserting a fragment of the *PHB3* promoter (2779 bp in front of ATG) upstream of *GUS*. The *p35s::IAA14-GFP* and *p35s::IAA28-GFP* constructs were generated by inserting fragments of the *IAA14* and *IAA28* open reading frames (ORFs) at the KpnI/SalI sites upstream of a *GFP* ORF previously inserted into pPZP211. *p35s::GATA23* and *p35s::LBD16* were generated by inserting the *GATA23* ORF and the *LBD16* ORF into the pPZP211 vector. The primer sequences are given in Supplemental Table 1. These constructs were transformed into *Agrobacterium tumefaciens* GV3101 and then into WT or *phb3ko* mutant plants by the floral-dip method. The transgenic plants were selected on 1/2 MS solid medium with 10 μg/mL kanamycin.

### Phenotypic analysis

*Arabidopsis* seeds were grown on 1/2 MS medium under long-day (LD; light: dark, 16 h: 8 h) conditions for 7 d, and then transferred into 1/2 MS medium added with 1 μM IAA, 100 μM SNAP, 500 μM cPTIO, or 1 μM NPA (or combinations thereof) for another 2 d. The numbers of LRs were then quantified and the length of PRs per seedling was measured under an Olympus BX53 fluorescence microscope. For gravitation assay, the seeds of WT and *phb3* mutants were grown vertically on 1/2 MS medium for 5 d and then rotated 90°. The LRP was observed under an Olympus BX53 fluorescence microscope 18 h later.

### GUS histochemistry

Histochemical staining for *pGATA23::GUS, pLBD16::GUS* and *pPHB3::GUS* activities in WT or *phb3* mutant plants was performed as previously described (Sessions *et al*., 1999) using 50 mM sodium phosphate buffer (pH 7.0) supplemented with 0.2% (v/v) Triton-X-100, 10 mM potassium ferrocyanide, 10 mM potassium ferricyanide, and 1 mM X-gluc. Samples were incubated overnight at 37□ and examined for GUS staining under a dissecting microscope (Xu *et al*., 2016).

### Endogenous NO detection in roots

The seedlings grown on 1/2 MS medium were washed twice with deionized water and incubated with 5 μM of the NO indicator DAF-FM DA for 15 min at room temperature in the dark. Then the seedlings were washed with fresh buffer (Tris-HCl, pH 7.0) for 15 min and endogenous NO was evaluated. DAF-FM DA reacts with NO to produce strong fluorescence with an excitation wavelength of 495 nm and emission wavelength of 515 nm. The endogenous NO was observed as green fluorescence under an Olympus BX53 fluorescence microscope.

### Gene expression analysis

Total RNA was extracted from the roots of seedlings previously treated with IAA or SNAP using an Ultrapure RNA Kit (CWBIO, China) (Cao *et al*., 2017). cDNA was then synthesized using EasyScript One-Step gDNA Removal and cDNA Synthesis SuperMix (TransGen, China). For qPCR, cDNA was conducted using a SYBR Green Master qPCR Kit (Roche Diagnostics, USA). The relative expression levels were calculated by the ΔΔC_T_ method (Schmittgen & Livak, 2008) using *ACT8* as the reference gene.

### Transcriptome analysis

*Arabidopsis* seeds were grown in 1/2 MS medium for 7 d. Total RNA of seedling roots was extracted using an Ultrapure RNA Kit, and the concentrations were detected using a NanoDrop 2000 spectrophotometer (Thermo). Three biological replicates were performed. The library was constructed and used for sequencing by Illumina HiSeq 2500 (GENE DENOVO, Guangzhou, China). For data processing, the clean reads were obtained from raw reads after filtering out adapters, poly(N), and low-quality reads. All the clean reads were then mapped to the *Arabidopsis* TAIR10.2 reference genome using TopHat (version 2.0.12). The fragments per kilobase of exon per million fragments mapped (FPKM) value was used to evaluate gene transcripts. *P* value < 0.01, false discovery rate (FDR) < 0.01, and fold change ≥ 2 were used as the principle (Li *et al*., 2017).

### Time-dependent fluorescence degradation assay

For fluorescence degradation assay, 7-day-old seedlings grown in 1/2 MS liquid medium were treated with 25 μM IAA and imaged with a Zeiss LSM880 (Zeiss) confocal laser-scanning microscope at 0, 5, 7, and 10 min after IAA treatment (Guseman *et al*., 2015).

## Results

### PHB3 is involved in LRP formation

To explore the molecular mechanism by which *PHB3* regulates LR development, two *phb3* mutants, *phb3-3* with a point mutation that converts Gly-37 to Asp and a T-DNA insertion line *phb3ko* (SALK_020707), were analyzed (Wang *et al*., 2010). The plants grew on 1/2 MS medium for 7 d followed by measurement of LR. The results showed that the LR density was significantly lower in *phb3* mutants (Figure 1A and 1B). We further determined the LRP number and found that the density of LRP was also obviously decreased in *phb3* mutants than in WT (Figure 1C), indicating that *PHB3* plays an important role in LRP formation.

**Figure 1.**
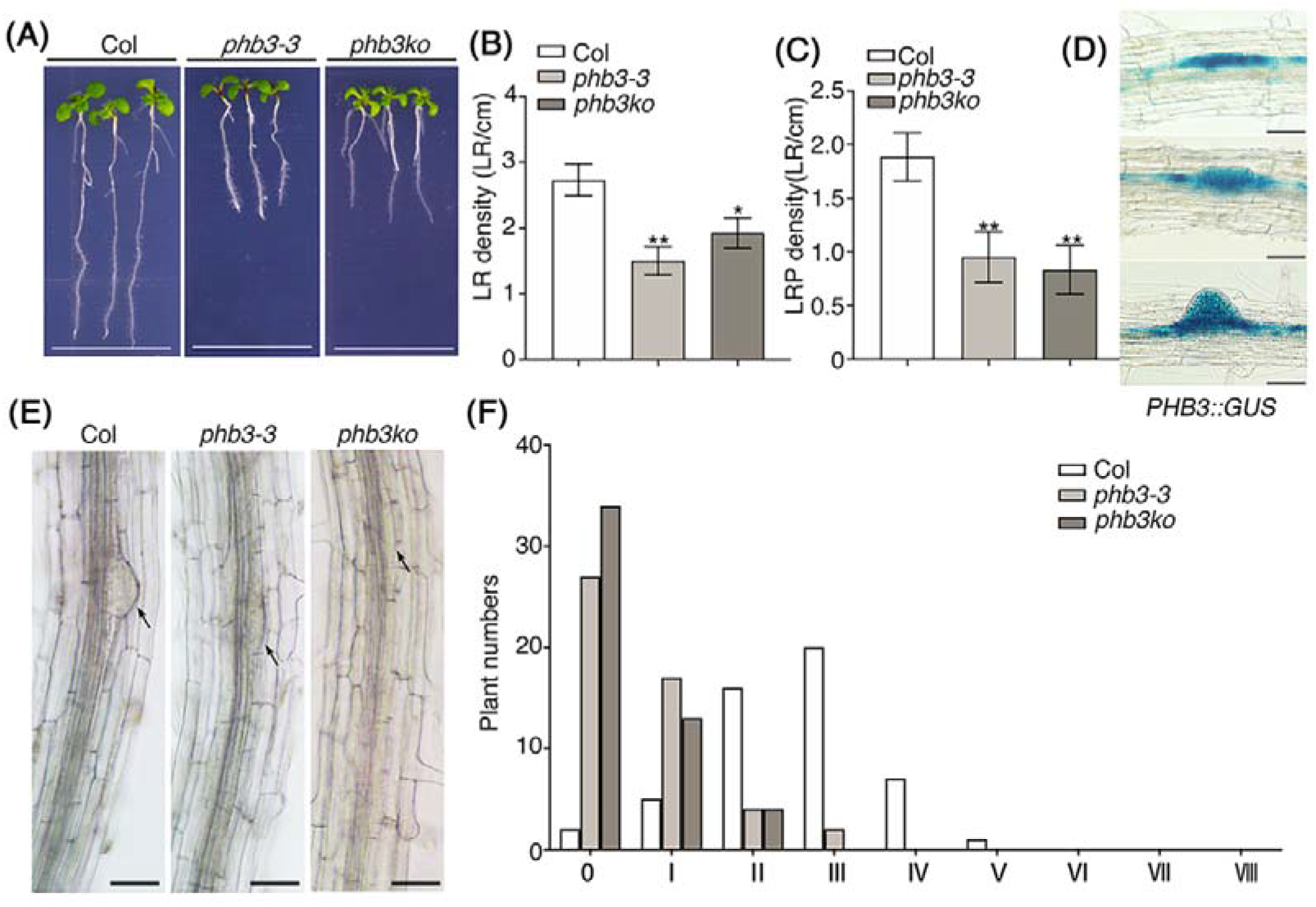
PHB3 is involved in LRP initiation. **(A)** LR growth in WT and *phb3* plants. Plants were grown on 1/2 MS medium for 7 d. Scale bars, 5 mm. **(B)** LR density in WT and *phb3* mutants. Plants were grown under the same conditions as in (A). LR density is the number of emerged LRs per 1 cm of the primary root length. Student’s *t*-test, \*\**P* < 0.01, \**P* < 0.05. The data are presented as means ± SD (n ≥ 20). **(C)** LRP density in WT and *phb3* plants. Plants were grown on 1/2 MS medium for 7 d. LRP density is the number of emerged LRs per 1 cm of the primary root length. Student’s *t*-test, ** *P* < 0.01. The data are presented as means ± SD (n ≥ 20). **(D)** The promoter activity of *PHB3* in the LRP different stages. Plants were grown on 1/2 MS solid medium for 7 d and GUS staining was detected. Scale bars, 50 μm. **(E)** LRP initiation in WT and *phb3* mutants. Plants were grown vertically on 1/2 MS medium for 7 d and then rotated 90°. LRP were counted using an Olympus BX53 fluorescence microscope after 18 h of rotation. The arrows represent LRP. Scale bars, 50 μm. **(F)** The quantitative results of (E), n = 50.

We also detected the expression of *PHB3* in WT roots using histochemical *β*-glucuronidase (GUS) analysis and found that *PHB3* was strongly expressed in the LRP (Figure 1D), consistent with its functions in LRP formation. To assess the LRP development of *phb3* mutants, we performed a gravitation assay, in which vertically cultured 5-day-old seedlings were rotated 90° and allowed to continue growing for 18 h before phenotypic observation. Normally, it results in increased auxin concentration and a new LRP to be initiated on the outside of the root curve (Laskowski *et al*., 2008; Guseman *et al*., 2015). Our results showed that the LRP appeared clearly at the curve site of WT roots but the emergence of LRP was obviously inhibited in the *phb3* mutants (Figure 1E). Furthermore, we counted the number of the LRP stages of the plants (n=50) at 18 h after rotation growth. The statistical data showed that the LRPs in WT were mainly at the second, third and fourth stages of LRP development while those in the *phb3* mutants were predominantly inhibited or at the first stage (Figure 1F). These results indicate that *PHB3* modulates the LRP formation.

### PHB3 regulates LRP initiation through NO

Our previous study has shown that both H2O2-induced NO accumulation and auxin-induced LR development are inhibited in *phb3* mutants (Wang *et al*., 2010). To investigate the molecular mechanism of NO participating in auxin-induced LR development, we analyzed the effects of NO on LR formation under our conditions. 5-day-old WT seedlings were treated with indole-3-acetic acid (IAA), auxin transport inhibitor naphthylphthalamic acid (NPA), NO donor S-nitroso-N-acetylpenicillamine (SNAP), and NO scavenger 2-(4-carboxyphenyl)-4, 4, 5, 5-tetramethylimidazoline-1-oxyl-3-oxide (cPTIO) for 2 d, and then assessed the LR density (Figure 2A and 2B). Exogenous IAA (1 μM) or SNAP (100 μM) treatments significantly increased the LR density of the seedlings, and application of both together had an even stronger effect, indicating that both IAA and NO promote LR formation under these conditions. Treatments with either NPA (1 μM) or cPTIO (500 μM) alone significantly reduced LR density and application of NPA and cPTIO together did not further enhance the reduction in LR density, suggesting that NO and IAA may function in the same pathway to regulate LR formation. Furthermore, treatment with SNAP restored LR formation in the NPA-treated seedlings while application of IAA did not rescue LR density in the cPTIO-treated seedlings, indicating that NO is required for auxin-induced LR development.

**Figure 2.**
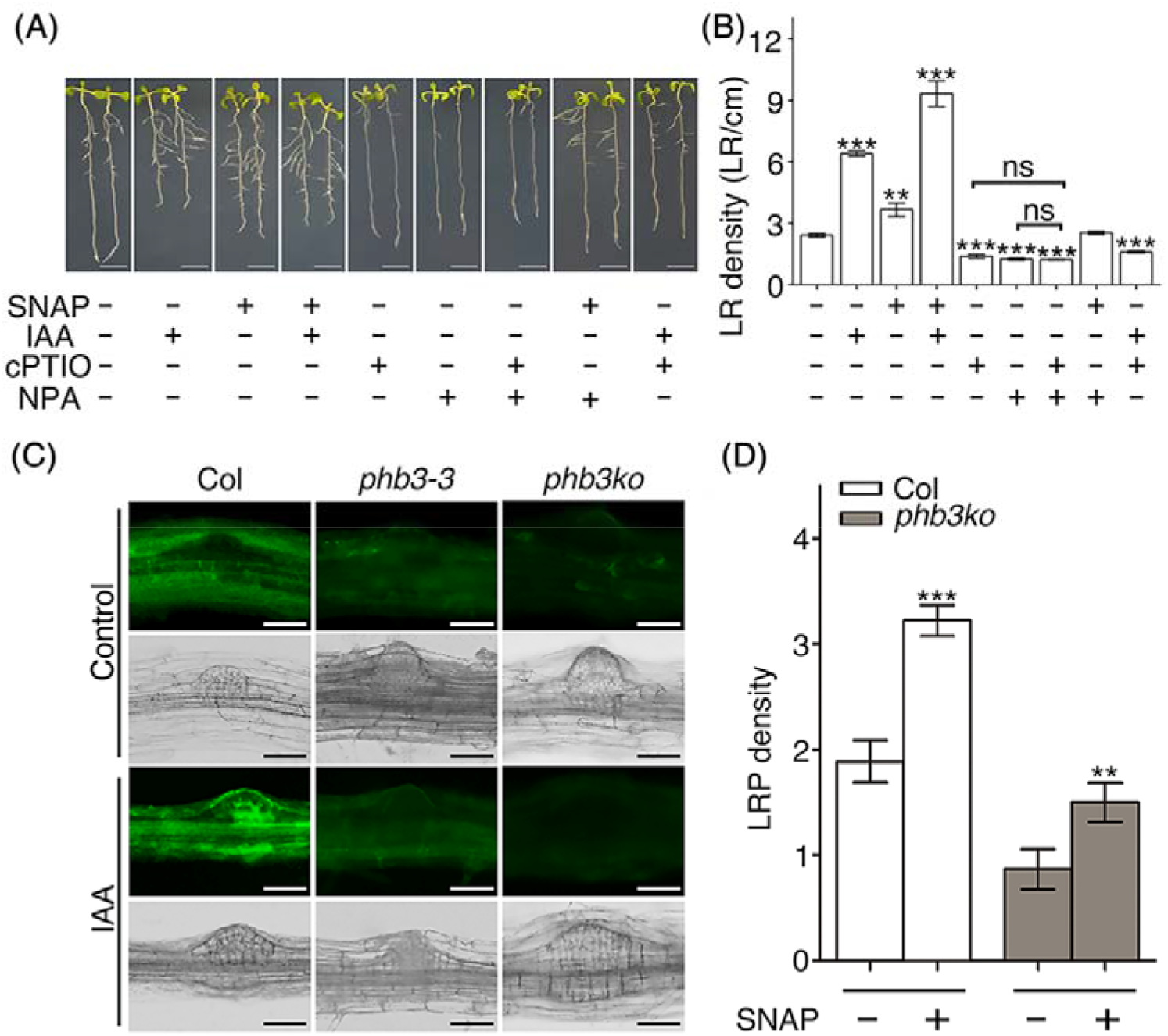
PHB3 regulates LRP initiation through NO. **(A)** WT plants were grown hydroponically in 1/2 MS medium for 5 d and then treated with various combinations of SNAP (100 μM), IAA (1 μM), cPTIO (500 μM), and NPA (1 μM) for 2 d. Scale bar, 5mm. **(B)** Quantitative data for LR density in WT plants after SNAP (100 μM), IAA (1 μM), cPTIO (500 μM), and NPA (1 μM) treatments. Student’s *t-*test, ns, not significant difference, ** *P* < 0.01, and *** *P* < 0.001. The data are presented as means ± SD (n ≥ 20). **(C)** NO accumulation in WT and *phb3* mutants. Plants were grown on 1/2 MS medium for 7 d. The seedlings were then washed twice with pure water and incubated with 5 μM DAF-FM DA for 15 min at room temperature in the dark. NO appeared as green fluorescence and was observed in primary roots (PR) and LRPs using an Olympus BX53 fluorescence microscope. Scale bars, 50 μm. **(D)** LRP density in NO-treated WT and *phb3ko* seedlings. Plants were grown on 1/2 MS medium for 5 d and then treated with 100 μM SNAP for 2 d. Student’s *t*-test, ** *P* < 0.01, *** *P* < 0.001. The data are presented as means ± SD (n ≥ 20).

As NO regulate auxin-induced LR development, and H_2_O_2_-induced NO accumulation in *phb3* mutants is reduced, it is possible that *PHB3* affects NO accumulation and thereby modulates LR development. To test this hypothesis, we further detected auxin-induced endogenous NO accumulation in the roots of WT and *phb3* plants using the NO indicator diaminofluorescein-FM diacetate (DAF-FM DA), which produces strong fluorescence after reacting with NO (Namin *et al*., 2013). The results showed that NO was mainly distributed around LRP, and the fluorescence of NO in the mutant was obviously weaker than that in WT (Figure 2C). After IAA treatment, the NO accumulation was strongly induced in the LRP of WT, while this induction was much lower in that of *phb3* mutants (Figure 2C). Additionally, application of NO donor SNAP increased LRP density of the *phb3* mutants (Figure 2D). These data suggest that *PHB3* participates in LRP development by mediating NO accumulation in plants.

To obtain molecular evidence of the effects of *PHB3* in regulating LR formation, we performed RNA sequencing (RNA-seq) analysis of WT and *phb3* plants grown on 1/2 MS medium for 7 d. The results showed 4,294 differentially expressed genes (DEGs) with 2,065 up-regulated and 2,229 down-regulated ones in *phb3ko* (Figure 3A and File S1). GO enrichment analysis for these DEGs revealed clusters related to root development, auxin biosynthesis and response, and NO response (Figure 3B and File S2). Three root development-related clusters are root morphogenesis, LR development, and post-embryonic root development. Three auxin-related clusters were enriched: response to auxin, auxin biosynthetic process, and auxin metabolic process. Two NO-related clusters were also found: response to NO and cellular response to NO. These results provide further evidence that *PHB3* is involved in regulating NO- and auxin-induced LR formation.

**Figure 3.**
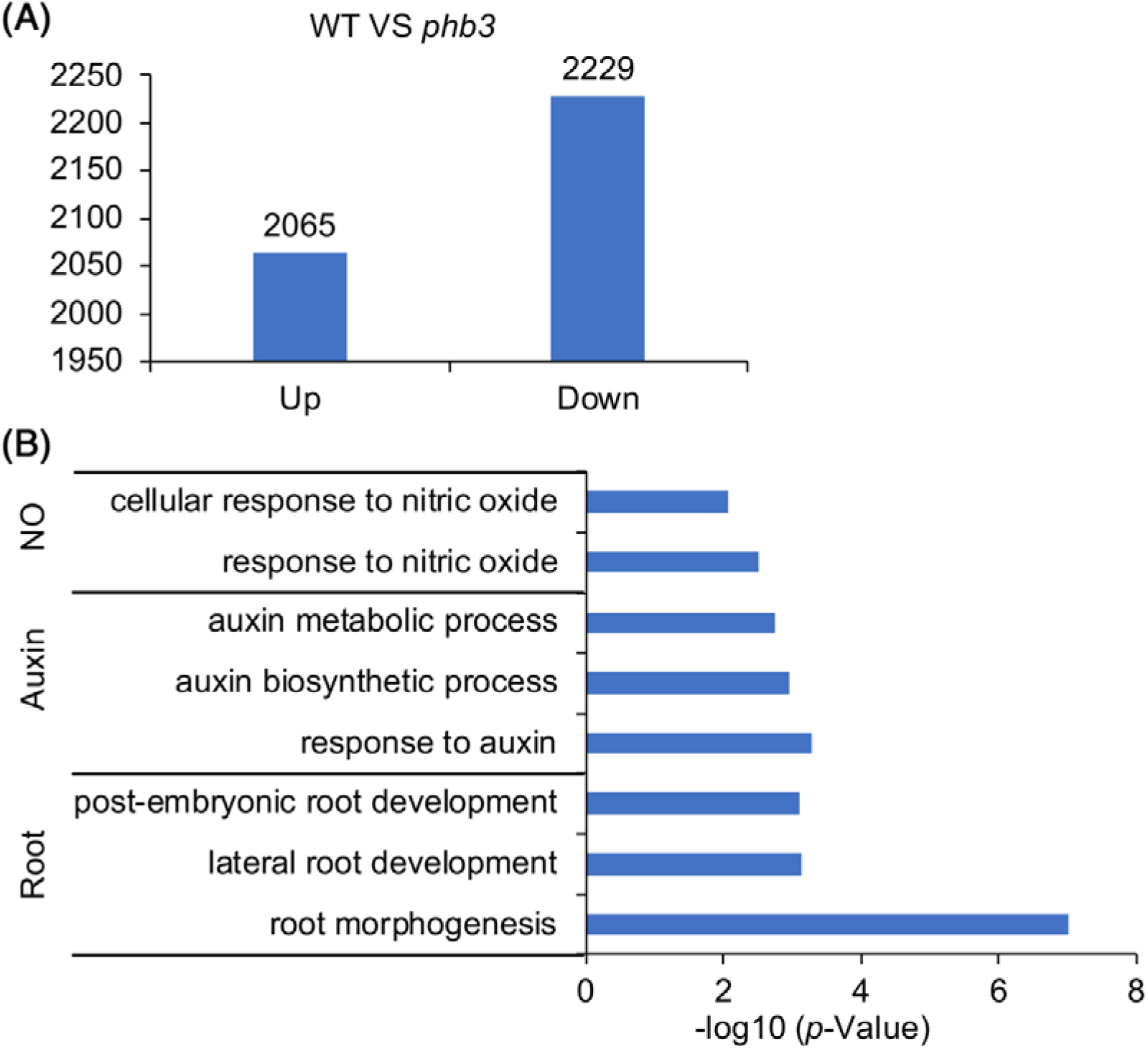
RNA-seq analysis for genes differentially expressed in WT and *phb3* plants. **(A)** Genes differentially expressed in WT and *phb3*. Plants were grown on 1/2 MS medium for 7 d, and then the roots were collected for RNA extraction used in RNA-s equencing. **(B)** GO analysis for genes differentially expressed in WT and *phb3* plants.

### *GATA23* and *LBD16* are involved in NO-induced LR development

To decipher further the mechanism by which NO regulates auxin-induced LR development, we investigated the transcript levels of four key genes involved in LRP formation (*GATA23, ARF19, LBD16*, and *LBD18*) in the roots of WT plants grown on 1/2 MS medium for 7 d. The results showed that the expression of *GATA23, ARF19, LBD16*, and *LBD18* was significantly induced by exogenous IAA (Figure 4A and Figure S1A), consistent with previous reports (Lavenus *et al*., 2013). After SNAP treatment, the expression of *GATA23* and *LBD16* was found to be increased rapidly (Figure 4B), but no significant change was found for *ARF19* and *LBD18* (Figure S1B). Consistent with these results, GUS-staining assay revealed that the activity of the *GATA23* and *LBD16* promoter was greatly enhanced by SNAP treatment (Figure 4C). These data suggest that the expression of *GATA23* and *LBD16* is strongly induced by NO.

**Figure 4.**
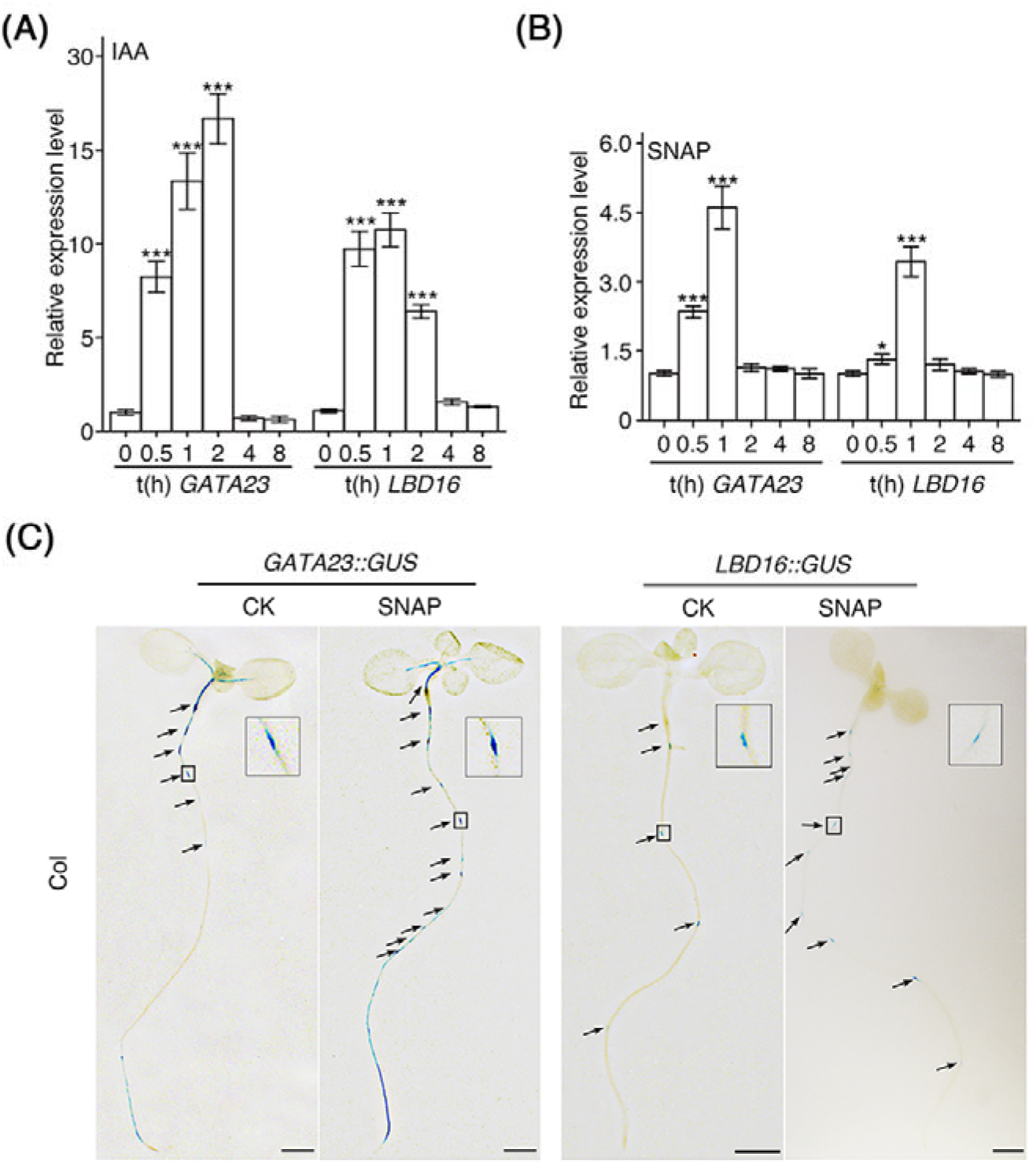
*GATA23* and *LBD16* are transcriptionally regulated by NO. **(A)** WT plants were grown on 1/2 MS medium for 7 d, and then treated with 1 μM IAA for 0 h, 0.5 h and 1 h, respectively, after which the roots were collected for RNA extraction. The expression of the *GATA23* and *LBD16* genes was determined by qPCR. Student’s *t*-test, *** *P* < 0.001. The data are presented as means ± SD (n ≥ 5). **(B)** WT plants were grown on 1/2 MS medium for 7 d and then treated with 100 μM SNAP for 0 h, 0.5 h and 1 h, respectively, after which the roots were collected for RNA extraction. The expression of *GATA23* and *LBD16* was determined by qPCR. Student’s *t*-test, * *P* < 0.05, *** *P* < 0.001. The data are presented as means ± SD (n ≥ 5). **(C)** Promoter activities of *GATA23* and *LBD16* in WT. Plants were grown on 1/2 MS medium for 7 d and then treated with 50 μM SNAP for 2 h. The promoter activities of the two genes are indicated by arrows. Scale bars, 50 μm.

To test the effects of NO on the expression of genes in plants, we performed RNA-seq using extracts of the roots of WT plants treated with SNAP. The results showed that the expression of 3,194 (1653+1541) genes was altered after SNAP treatment (Figure 5A, File S3), among which many LR-regulated genes, including IAA5, ILL6, PIN6, AIR12, IAA1, ARF16, PIN5, GATA23 and ILL3, showed up-regulation (Figure 5B). GO analysis revealed that three auxin-related clusters were enriched among these genes: response to auxin, auxin metabolic process, and auxin biosynthetic process (Figure 5C, File S4). The above data suggest that NO may play important roles in auxin signaling and metabolism.

**Figure 5.**
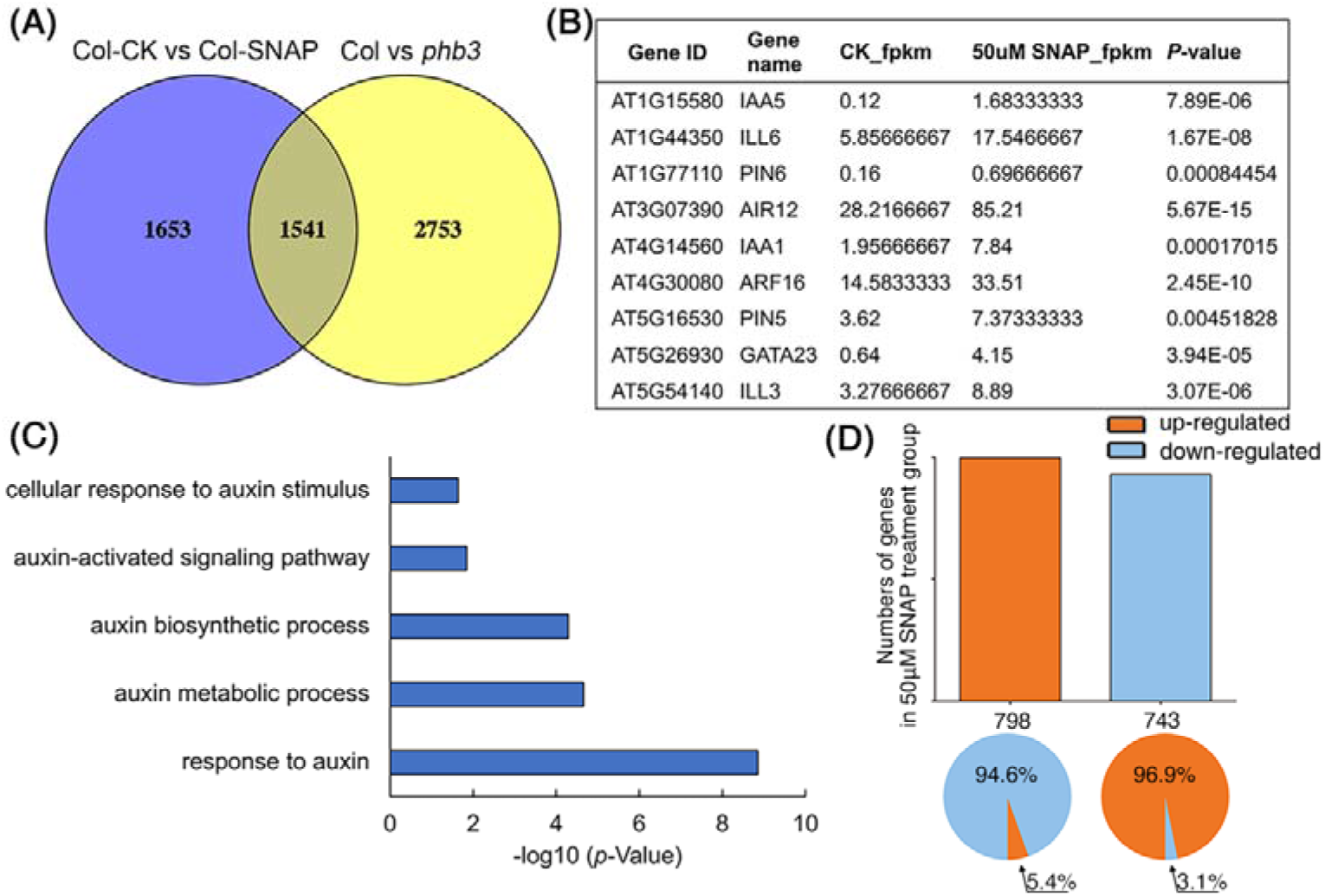
Analysis for DEGs in WT after NO treatments and DEGs in *phb3* mutant. **(A)** Venn diagram showing the number of genes differentially expressed in *phb3* mutant, and in WT roots by treatments with different concentrations of SNAP (50 μM) for 1 h compared with control plants (0 μM SNAP). Plants were grown on 1/2 MS medium for 7 d, and then treated with different concentrations of SNAP (0 μM and 50 μM) for 1 h, after which the roots were collected for RNA extraction used in RNA-sequencing. **(B)** Known LR-regulated genes with DEGs in in WT after SNAP treatment. **(C)** GO analysis for DEGs in WT after SNAP treatment. **(D)** Transcriptome analysis of WT and *phb3* vs the SNAP-treated WT.

We compared the DEGs between WT and *phb3* vs the SNAP-treated WT (Figure 5A), and found that 1541 DEGs in *phb3* mutant was regulated by NO, which is about 35.9% (1541/4294) of total DEGs in *phb3* mutant. Furthermore, 798 DEGs were up-regulated by SNAP, while 94.6% (755/798) of 798 DEGs were down-regulated in *phb3* mutant (Figure 5D). Meanwhile 743 were down-regulated by SNAP treatment, among which 96.9% (720/743) were up-regulated in *phb3* mutant (Figure 5D). These data indicate that most of genes up-regulated in *phb3* mutant are down-regulated in SNAP-treated WT and vice versa. This opposite DEGs between SNAP treatment and *phb3* mutant strongly suggest an important link between the function of PHB3 and NO signaling.

Previous studies have shown that both *GATA23* and *LBD16* regulate LR formation (De Rybel *et al*., 2010; Goh *et al*., 2012). Consistent with these findings, we also found that LR development was inhibited in *gata23* and *lbd16* mutants (Figure S2). To examine the roles of *GATA23* and *LBD16* in NO-induced LR development, we investigated the effects of exogenous SNAP treatment on WT, *gata23*, and *lbd16* plants. After a 2-d treatment with SNAP (100 μM), the LR density was strongly induced and increased 282% that of untreated control in the WT, but only increased 43% in *gata23* and 149% in *lbd16*, respectively (Figure 6A and 6B). These results indicate that both *GATA23* and *LBD16* contribute to NO-induced LR formation.

**Figure 6.**
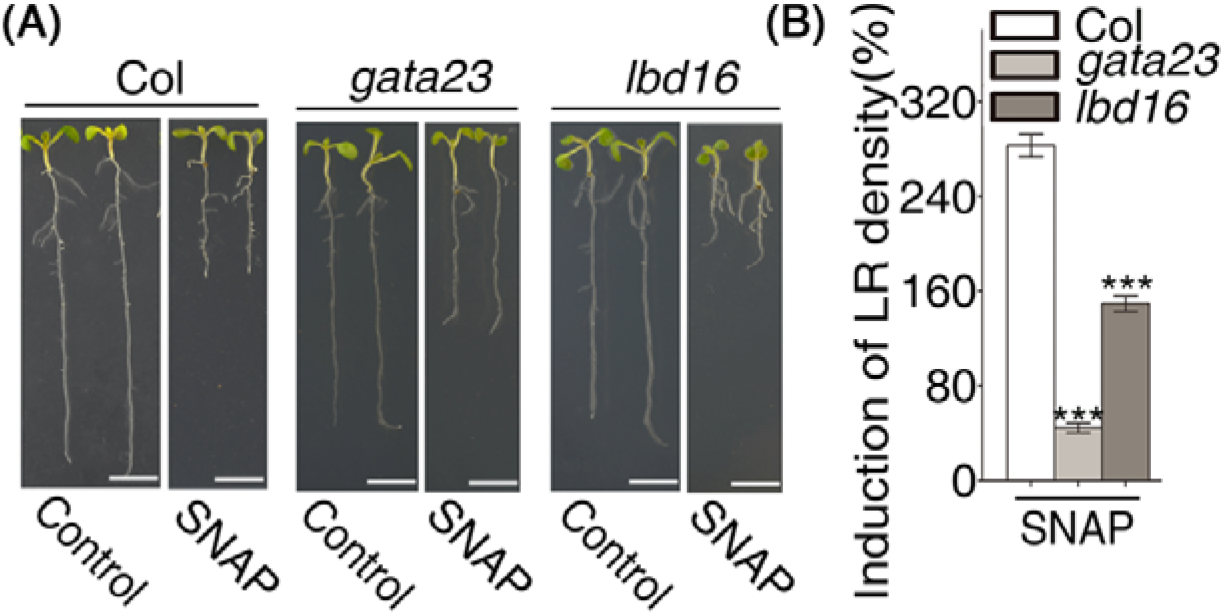
*GATA23* and *LBD16* are involved in NO-induced LR development. **(A)** WT, *gata23*, and *lbd16* plants were grown on 1/2 MS liquid medium for 5 d, and then treated with SNAP (100 μM) for 2 d. Scale bars, 5 mm. **(B)** Density of LRs was quantified by microscopy observation. Student’s *t*-test, *** *P* < 0.001. The data are presented as means ± SD (n ≥ 20).

### PHB3 regulates the expression of *GATA23* and *LBD16* by modulating NO-mediated AUX/IAA degradation

Considering that the accumulation of NO is strongly reduced in *phb3* mutants and that the expression of LR regulatory genes (*GATA23* and *LBD16*) is enhanced by NO, we speculated that *PHB3* may influence LRP development by modulating the accumulation of NO. Therefore, we detected the expression of *GATA23* and *LBD16* by real-time qPCR. The results showed that the expression of *GATA23* and *LBD16* was significantly reduced in *phb3* plants (Figure 7A).

**Figure 7.**
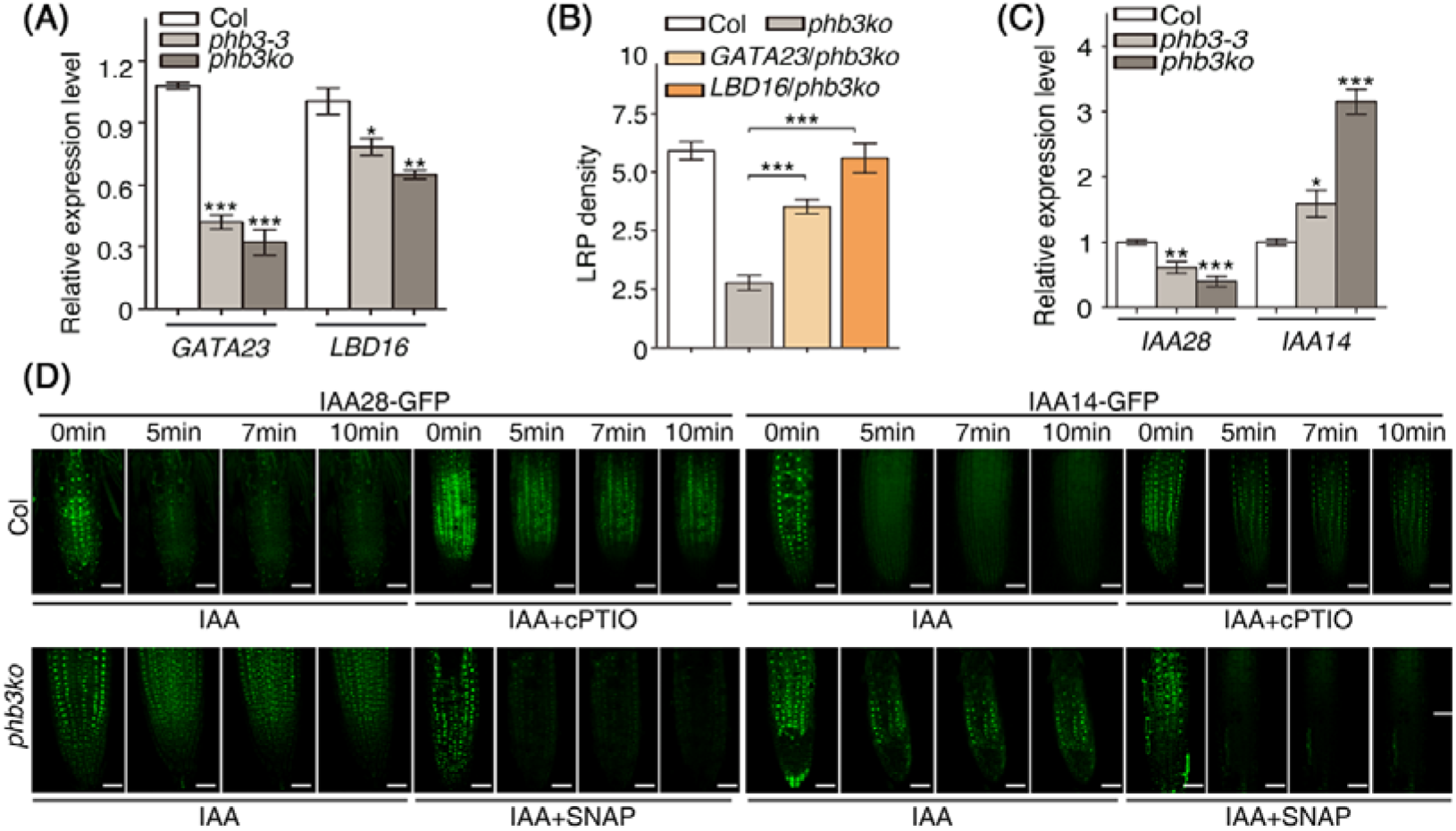
*PHB3* affects the expression of *GATA23* and *LBD16* through NO-mediated degradation of IAA14/28. **(A)** Expression of *GATA23* and *LBD16* in WT and *phb3* mutants. Plants were grown on 1/2 MS medium for 7 d and then roots were collected for RNA extraction. The expression of the genes was determined by qPCR. Student’s *t*-test, * *P* < 0.05, \*\**P* < 0.01, *** *P* < 0.001. The data are presented as means ± SD (n ࣙ 5). **(B)** LRP density in WT, *GATA23/phb3ko*, and *LBD16/phb3ko* plants. Plants were grown on 1/2 MS medium for 7 d. Student’s *t*-test, *** *P* < 0.001. Data are presented as means ± SD (n ≥ 20). **(C)** Expression of *IAA28* and *IAA14* in WT and *phb3* mutants. Plants were grown on 1/2 MS medium for 7 d and then roots were collected for RNA extraction. The expression of the two genes was determined by qPCR. Student’s *t*-test, * *P* < 0.05, ** *P* < 0.01, *** *P* < 0.001. The data are presented as means ± SD (n ≥ 5). **(D)** Degradation of IAA28-GFP and IAA14-GFP in the roots of WT and *phb3ko* mutant. Plants were grown on 1/2 MS medium for 7 d and then treated with 25 μM IAA, 25 μM IAA +500 μM cPTIO, and 25 μM IAA +100 μM SNAP, respectively. The green fluorescence of IAA-GFP was observed with a Zeiss LSCM high-resolution laser confocal microscope. Scale bars, 500 μm.

To determine whether *GATA23* and *LBD16* are functionally required for LRP initiation, we ectopically expressed *GATA23* and *LBD16* separately in the *phb3ko* mutant. The results showed increased the LRP density in transgenic plants of *GATA23* and *LBD16* genes (Figure 7B), indicating that both genes work downstream of *PHB3* to modulate LRP formation.

The auxin regulatory protein IAA28 plays a key role in specifying pericycle cells for development into LR founder cells prior to LR initiation (De Rybel *et al*., 2010) and another protein, IAA14, is involved in the asymmetric division of LR founder cells that occurs as a part of LR initiation (Fukaki *et al*., 2005; Goh *et al*., 2012). Therefore, we investigated the transcript levels of *IAA28* and *IAA14* in *phb3* mutants. Interestingly, the expression of *IAA28* was significantly decreased and that of *IAA14* was increased in the mutants (Figure 7C), indicating that *PHB3* modulates the transcript levels of *IAA28* and *IAA14*. In theory, the decreased expression of *IAA28* should result in increased expression of *GATA23* (De Rybel *et al*., 2010), which is the opposite of the reduced expression of *GATA23* found in *phb3* mutants. So, we further performed a protein degradation assay using fluorescent IAA-GFP protein, which has been widely used as an in *vivo* marker to detect the degradation of SCF^TIR1^-AUX/IAA in the nucleus (Guseman *et al*., 2016). In our assay, we transformed WT and *phb3* plants with the constructs *IAA28-GFP* and *IAA14-GFP*, respectively. The transgenic plants were grown on 1/2 MS solid medium for 7 d, and then treated with 25 μM IAA. Fluorescence microscopy assessment showed that the IAA28-GFP and IAA14-GFP signals disappeared within 5 min of IAA treatment in the root cells of WT, but persisted in the root cells of *phb3* plants even after 10 min of IAA treatment (Figure 7D). In addition, we treated the *phb3* mutant with SNAP and found that the degradation of IAA28 and IAA14 was restored after NO supply (Figure 7D). When treated with cPTIO, WT plants showed inhibited degradation of IAA28 and IAA14 proteins (Figure 7D). These data indicate that NO plays a key role for the degradation of IAA28 and IAA14 and that PHB3 regulates LRP initiation by modulating NO-mediated AUX/IAA degradation.

## Discussion

In nature, plants constantly adapt to the changing environment by adjusting their root architecture. The LR is a critical factor determining root architecture (Nibau *et al*., 2008). Understanding the regulatory mechanisms of LR development is thus of great importance in efforts to enhance the yield and stress resistance of crops. Auxin plays an essential role in LR development, including LRP initiation, LRP formation, and LR growth (Fukaki *et al*., 2005; De Rybel *et al*., 2010; Lee *et al*., 2013; Jeon *et al*., 2017). NO, as an important signaling molecule plays a crucial role in many plant developmental processes including root development. NO has been found to regulate auxin-induced LR development in tomato (Correa-Aragunde, Graziano, & Lamattina, 2004) and to promote the S-nitrosylation of TIR1, which enhances the interaction of TIR1/AFB with AUX/IAA, resulting in the expression of IAA-induced genes (Terrile *et al*., 2012).

We previously showed that H_2_O_2_-induced NO accumulation is diminished and the number of LRs is much lower in *phb3* mutants after auxin treatment (Wang *et al*., 2010). However, the underlying mechanism whereby PHB3 regulates LR development remained unclear. Here, we found that the density of LRPs was reduced in *phb3* mutants, indicating that *PHB3* is required in LRP formation (Figure 1A-1C). Furthermore, our data showed reduced number and the growth rate of LRPs in *phb3* mutants (Figure 1A-F), indicating that *PHB3* regulates LR formation by influencing the number and growth rate of LRPs.

Dr. Lamattina’s group reported auxin-induced NO accumulation in tomato pericycle cells undergoing anticlinal or periclinal division (Correa-Aragunde, Graziano, & Lamattina, 2004), suggesting that NO plays an important role in the early development of LRP. Application of the cPTIO inhibited auxin-induced LR formation. When tomato plants were treated with NPA, NO treatment could still promote LR initiation in the roots (Correa-Aragunde, Graziano, & Lamattina, 2004). We investigated the effects of NO on auxin-induced LR formation using different combination treatments of IAA, IAA transport inhibitor, NO donor, and NO scavenger (Figure 2A-B). Our data indicate that NO plays an essential role in IAA-induced LR development, which is consistent with the previous results from tomato (Correa-Aragunde, Graziano, & Lamattina, 2004).

However, the mechanism by which PHB3 contributes to NO accumulation remains unknown. It has been demonstrated that PHB3 is dually localized in the mitochondria and chloroplasts of plant cells (Ahn *et al*., 2006; Van Aken *et al*., 2007; Seguel *et al*., 2018). Both hydrogen peroxide (H_2_O_2_) and superoxide ion 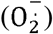 have been found to accumulate in the mitochondria of *phb3* mutant cells (Kong *et al*., 2018), indicating that PHB3 is involved in regulating the homeostasis of reactive oxygen species. Consistently, we performed RNA-sequencing on *phb3* mutant and GO analysis revealed two hydrogen peroxide-related clusters: regulation of hydrogen peroxide metabolic processes and hydrogen peroxide biosynthesis. In addition, NO has been shown to be consumed by reacting with 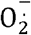 in potato tuber mitochondria (de Oliveira *et al*., 2008). In animals, excess 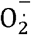 can convert NO into ONOO^-^ and thereby decrease NO bioavailability (Brown & Borutaite, 2007; Schiffrin, 2008; Brandes *et al*., 2011). We also found that the endogenous NO content was reduced in the roots of *phb3* mutants (Figure 2C). These findings implicate that the excess H_2_O_2_ and 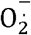 may convert NO to some intermediates, leading to the reduced accumulation of endogenous NO in *phb3* mutants.

In the efforts to decipher the mechanism of NO in regulating LR formation, we found that the expression of the root regulatory genes *GATA23* and *LBD16* can be induced by NO (Figure 4), suggest that NO regulates the LR development by promoting the specification of founder cells and the asymmetric division of founder cells. Further investigation revealed that the expression levels of these two genes were significantly decreased in the *phb3* mutants (Figure 7A), suggesting that *PHB3* may affect LRP formation by regulating the expression of these two genes. In addition, the LRP density of the *phb3* mutant could be rescued by over-expression of *GATA23* or *LBD16* (Figure 7B), indicating that *GATA23* and *LBD16* act downstream of *PHB3*. These results suggest that PHB3 may regulate LR development by affecting the accumulation of NO and then modulating the expression *GATA23* and *LBD16*.

As the expression of *GATA23* and *LBD16* is regulated by IAA28 and IAA14, respectively (Lavenus *et al*., 2013), we investigated *IAA28* and *IAA14* expression in the *phb3* mutants. The expression of *IAA28* was inhibited while the expression of *IAA14* was increased in *phb3* mutants (Figure 7C). Decreased expression of *IAA28* should result in increased expression of *GATA23*. But actually, reduced expression of *GATA23* was found in *phb3* mutants. These results implicate that the regulation of *GATA23* by IAA28 may not at the transcriptional levels but at protein levels. In addition, NO can promote the degradation of AUX/IAA proteins through enhancing the interaction between TIR1/AFB and AUX/IAA (Terrile *et al*., 2012). However, the mechanism by which AUX/IAA proteins participate in NO-regulated LR development was still unclear. Therefore, we performed AUX/IAA protein degradation assays and found that IAA28-GFP and IAA14-GFP degradation was strongly delayed in *phb3* mutant after IAA treatment (Figure 7D). Furthermore, the degradation of both proteins was restored in *phb3* mutant after NO treatment. Thus, the decreased expression of *GATA23* is caused by NO-mediated delay of IAA28 protein degradation and not by decreased gene expression of *IAA28*. Similarly, the decreased gene expression of *LBD16* results from delayed degradation of IAA14 protein.

Taken together, our findings suggest that *PHB3* regulates the degradation of IAA28 and IAA14 by controlling endogenous NO concentrations, thereby regulating the expression of downstream genes (*GATA23* and *LBD16*) and ultimately modulating the founder cell identity and asymmetric division (Figure 8). The *PHB3-*mediated regulation of AUX/IAA proteins and the downstream genes *GATA23* and *LBD16* provides insight into the cross-talk between NO and auxin-induced LR formation and a new mechanism of IAA-mediated LR development.

**Figure 8.**
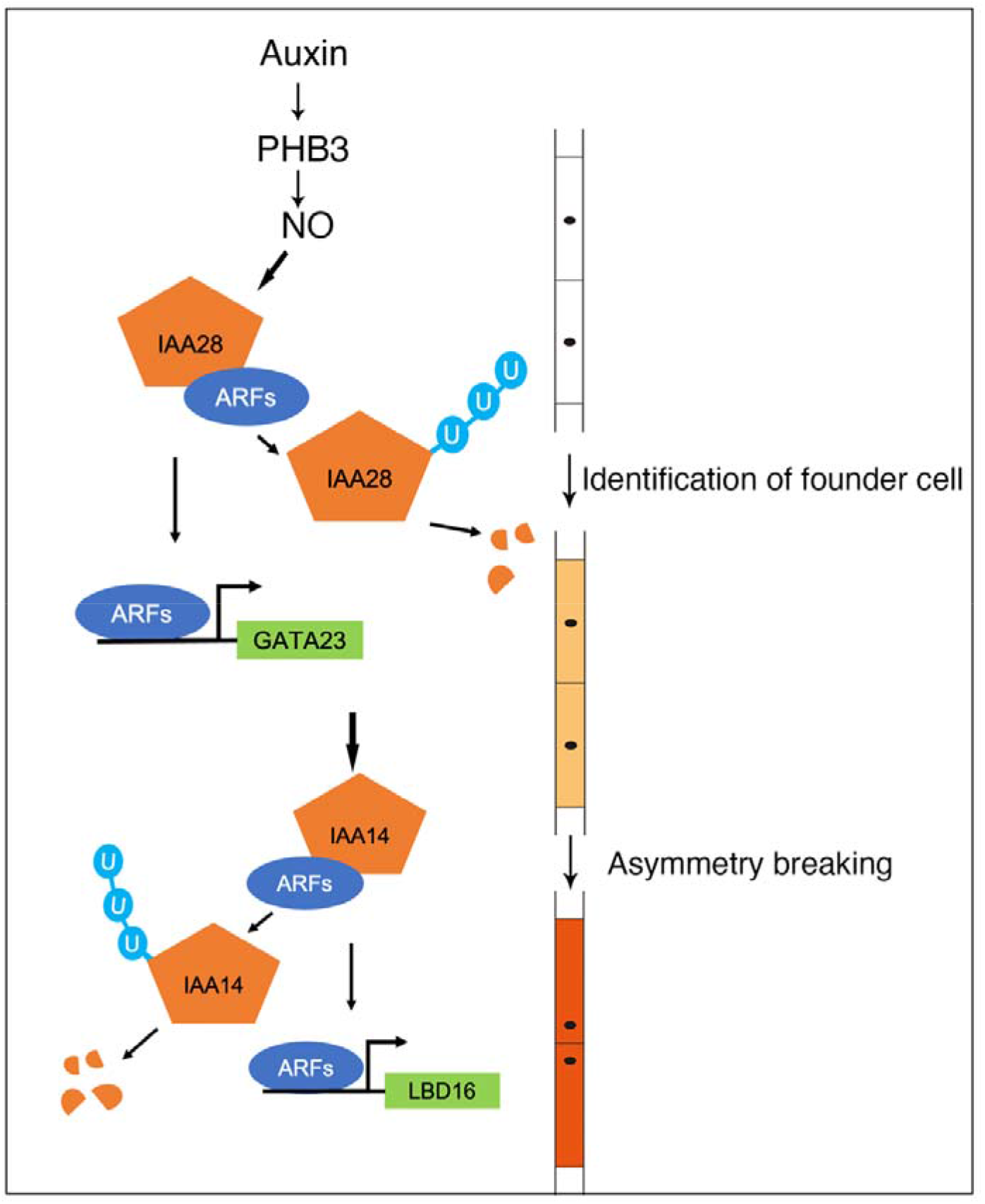
Model showing the roles of *PHB3* in regulating LRP initiation. PHB3 regulates LRP initiation by controlling the levels of NO. NO may promote the IAA-mediated degradation of IAA28 and release of ARFs. Released ARFs can bind to the promoter and activate the expression of downstream gene *GATA23* to control LR founder cell specification. Then NO may promote the degradation of IAA14 and release of its interacted ARFs. The released ARFs can bind to the promoter and activate the expression of the *LBD16* to initiate asymmetric division of LR founder cells.

## Supplementary data

**Supplemental Table S1.** Primer list

**Supplemental Figure S1.** The expression of the LR regulatory genes *ARF19* and *LBD18* after IAA and SNAP treatment.

**Supplemental Figure S2.** LR development is inhibited in *gata23* and *lbd16* mutants.

**Supplemental File S1.** Genes whose expression changed more than 2-fold in WT and *phb3* mutant.

**Supplemental File S2.** Some clusters for genes differentially expressed in WT and *phb3* mutant.

**Supplemental File S3.** Genes whose expression changed more than 2-fold after NO treatment.

**Supplemental File S4.** Some clusters for genes differentially expressed in WT and SNAP-treated.

## Acknowledgements

We thank Prof. Tom Beeckman for providing *GATA23::GUS* transgenic *Arabidopsis* lines. This work was supported by NSFC (31970270) and Funds of Shandong “Double Tops” Program to Y.W.

## Author Contributions

Y.W. designed the research. S.L., Q.L., X.T, L.M, M.L., X.W., N.L., F.L., and J.S. performed the experiments. Q.L., Y.W., and S.L. analyzed the data. Y.W., N.M.C, and Q.L. wrote the manuscript.

